# NAStructuralDB : Structural database to facilitate computational studies of molecular modeling and recognition of proteins with special focus on antibody-antigen interactions

**DOI:** 10.1101/2025.10.17.683056

**Authors:** Dawid Chomicz, Paweł Dudzic, Sonia Wrobel, Tomasz Gawłowski, Samuel Demharter, Roberto Spreafico, Hervé Minoux, Andrew Phillips, Konrad Krawczyk

## Abstract

Studying the interactions between antibodies and antigens is fundamental to the development of novel therapeutic biologics. Predictions of such interactions start with data collection. Though there exist reliable resources to identify antibody structures in the Protein Data Bank (PDB), such data still requires substantial processing to be usable in predictive tasks. Redundancy in sequences needs to be removed to avoid data leakages between train, test and validation sets. Descriptors such as surface accessibility, secondary structure and antibody region information need to be additionally annotated. Information on inter- and intra-molecular contacts, which is crucial to studying paratope/epitope information, needs to be collected. The specialized immunoglobulin format of Nanobodies® requires a separate dataset mirroring that of antibodies, given that their structure contains only a single VHH chain. Because antibody-antigen structures account for a small amount of all protein-protein contacts, having a molecular contact reference from other proteins is also desired. To address these issues, we introduce NAStructuralDB (https://naturalantibody.com/na-structural/), a dataset of processed structures of antibodies, Nanobodies®, proteins and their complexes with molecular contact information and associated annotations. We use the opportunity of having collected the contact data to provide a reference of binding propensities of different residues across distinct contact types. We anticipate that this dataset will accelerate a broad range of predictive tasks by standardizing common, time-consuming data preparation steps in antibody and protein design.

## Introduction

Antibody-antigen binding is a subproblem of a more generic protein-protein binding problem (Esmaielbeiki et al. 2016). Despite progress with protein structure prediction (Jumper et al. 2021), predicting the structure of protein assemblies and complexes still remains a challenge (Kryshtafovych et al. 2021; Evans et al. 2022; Wong et al. 2022). Discerning protein-protein specificity has long been a goal of protein design (Watson et al. 2022), a solution to which could radically accelerate our ability to discover novel therapeutics. This is particularly desired for antibodies, which are the top class of biologics and consistently account for approximately half of the top ten blockbuster drugs in the United States by sales (Martin et al. 2023; Lu et al. 2020).

Antibodies have different molecular contact preferences than ‘general’ proteins (Krawczyk et al. 2013), requiring the development of a specific set of binding prediction tools (Norman et al. 2020; Wilman et al. 2022). There are several classes of antibody-specific binding predictors that can be distinguished based on whether they predict the binding site on individual molecules or on their pairs. Paratope predictors focus on annotating the antibody-binding site, which is typically a subset of the Complementarity Determining Regions (CDRs). Structural approaches have been proposed (Krawczyk et al. 2013; Chinery et al. 2023), however competitive performance can be achieved with sequence alone in the absence of antigen information (Liberis et al. 2018; Leem et al. 2022). Epitope prediction aims to detect a region on an antigen that could be an antibody target, where discontinuous epitopes typically form the majority of antibody-recognized sites. Here, multiple approaches perform predictions based on structure, without taking the partner antibody into account (Tubiana,

Schneidman-Duhovny, and Wolfson 2022). However, including antibody information has been shown to provide more information than the antigen alone for epitope prediction (Jespersen et al. 2019; Krawczyk et al. 2014). Taking both antibody and antigen binding into account and predicting the structure of their complex has long been a major problem in protein-protein docking (Sircar and Gray 2010; Ambrosetti et al. 2020). Antibody-specific methods are necessary in this area, however there is still no generalizable solution that can reconstruct complexes from separated antibody and antigen structures, with the performance dropping significantly for structure prediction models (Schneider et al. 2021; Jin, Barzilay, and Jaakkola 17--23 Jul 2022; Sircar and Gray 2010; Krawczyk et al. 2014). One can also attempt antibody design, wherein a binder is created for a specific epitope, by traditional structural methods or protocols using novel generative concepts (Adolf-Bryfogle et al. 2018; Rangel et al. 2022; Jin, Barzilay, and Jaakkola 17--23 Jul 2022).

Data preparation is arguably the most important and time-consuming step when creating input for model training. Structural antibody data is deposited in the Protein Data Bank (Berman et al. 2000), while specialized sources such as ABDB (Ferdous and Martin 2018) and the Structural Antibody Database (Dunbar et al. 2014) filter out the antibodies from the larger PDB repository. While these databases allow identifying antibodies and filtering by antigen, further processing steps involving epitope/paratope identification are not performed. This gap motivates the creation of solutions tailored to antibody-antigen interactions (Huang et al. 2025; Zhou et al. 2024).

Since further processing of structures is needed for countless downstream applications, various software solutions and websites have been created to address this need. DeepRank (Renaud et al. 2021) was created to help process the structural information into structural formats useful for training interface predictors. Such frameworks process structures into feature-rich datasets, extracting labels such as number of residue-residue contacts, van der Waals energy, electrostatic energy and more. Such inputs can then be used within the same framework to train accurate rescoring of docking poses based on physicochemical descriptors. Another example of a framework that features protein-protein contacts is Arpeggio (Jubb et al. 2017). Upon structure submission, interface interactions are labeled and grouped by hydrogen bonds, van der Waals or ionic interactions. Websites such as Protein Contact Atlas offer processing and visualization of non-covalent contacts available in the Protein Data Bank (Kayikci et al. 2018).

Though such frameworks offer ways to process and visualize molecular contacts, they rarely offer bulk downloads of processed data from the Protein Data Bank, nor do they focus on antibodies. In order to provide a standardized shortcut from data acquisition to model training, we created the NaturalAntibody Structural Database (NAStructuralDB). The database consists of general proteins, antibodies, Nanobodies®, and their interfaces, ready for deep learning featurization on par with state of the art use-cases for structure predictions (Eshak and Goupil-Lamy 2025; Hitawala and Gray 2024), co-folding (Boitreaud et al. 2024; Wohlwend et al. 2024), diffusive design (Bennett et al. 2024), docking (Ambrosetti et al. 2020; Sircar and Gray 2010; Jeliazkov et al. 2021) or binding prediction (Jespersen et al. 2019; Liberis et al. 2018). We anticipate that our resource will provide a standardized dataset to facilitate reproducibility and save time in training machine learning models for antibody discovery.

## METHODS

### Database Contents

*Overview*.

NAStructuralDB is a database of processed structures derived from the Protein Data Bank. To contextualize its features and capabilities among other available resources, a detailed side-by-side comparison with other major structural interaction databases is provided in Table 1.

**Table 1.**
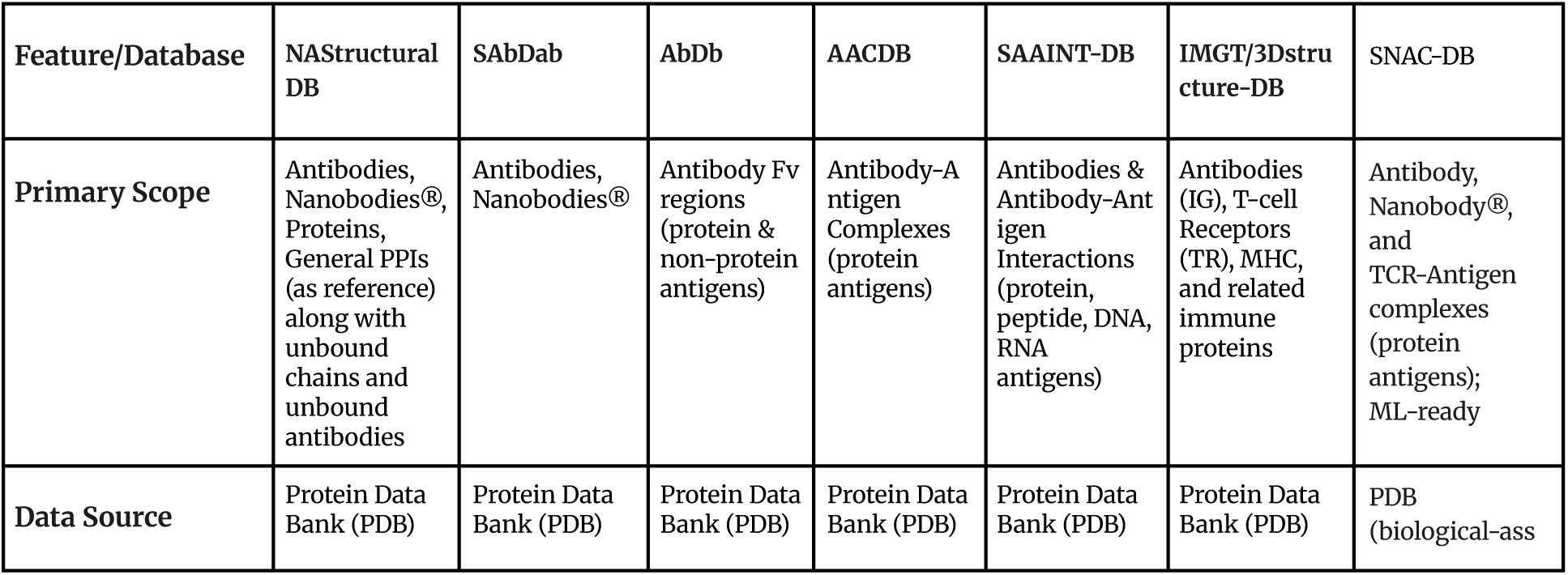
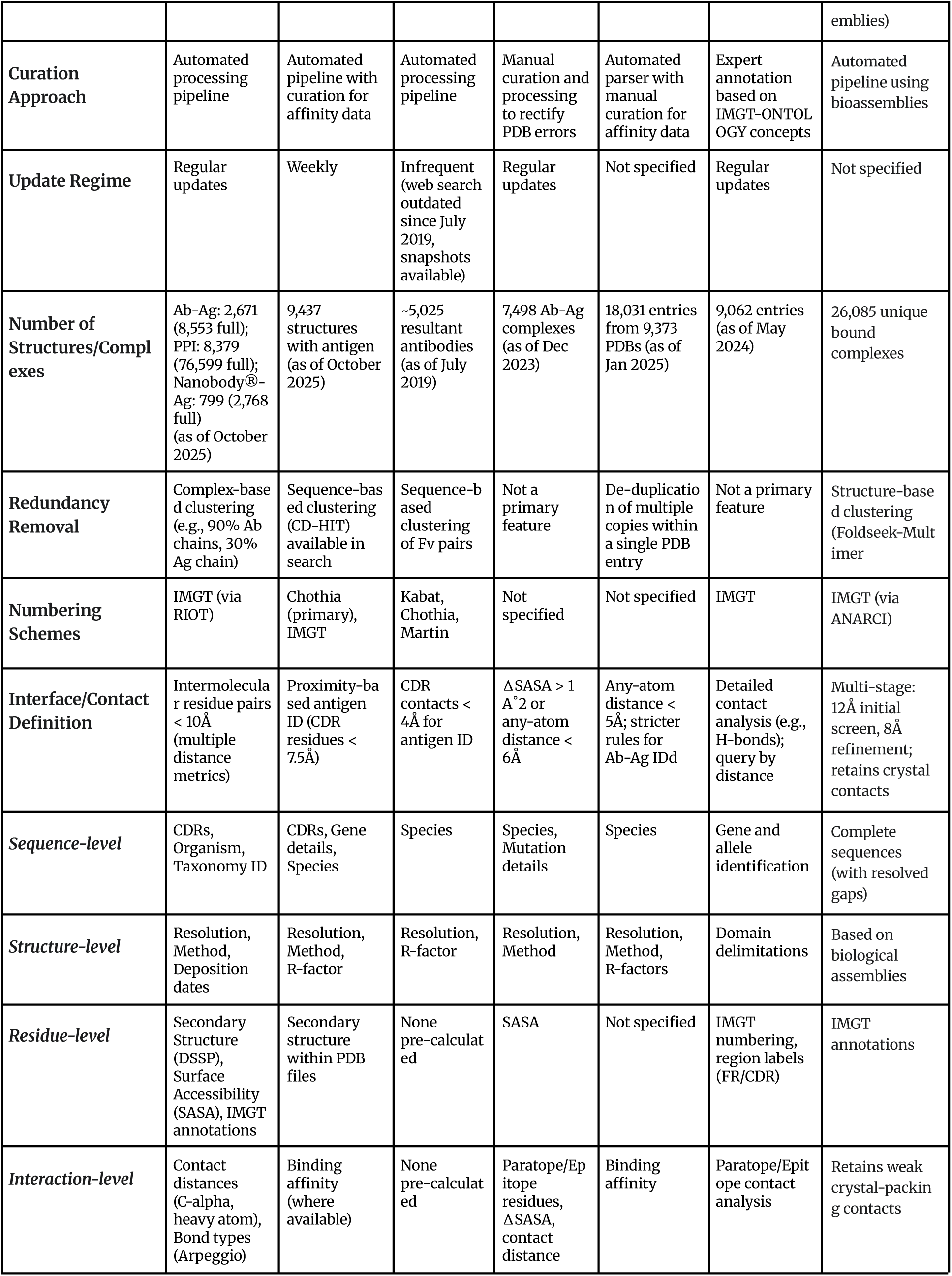
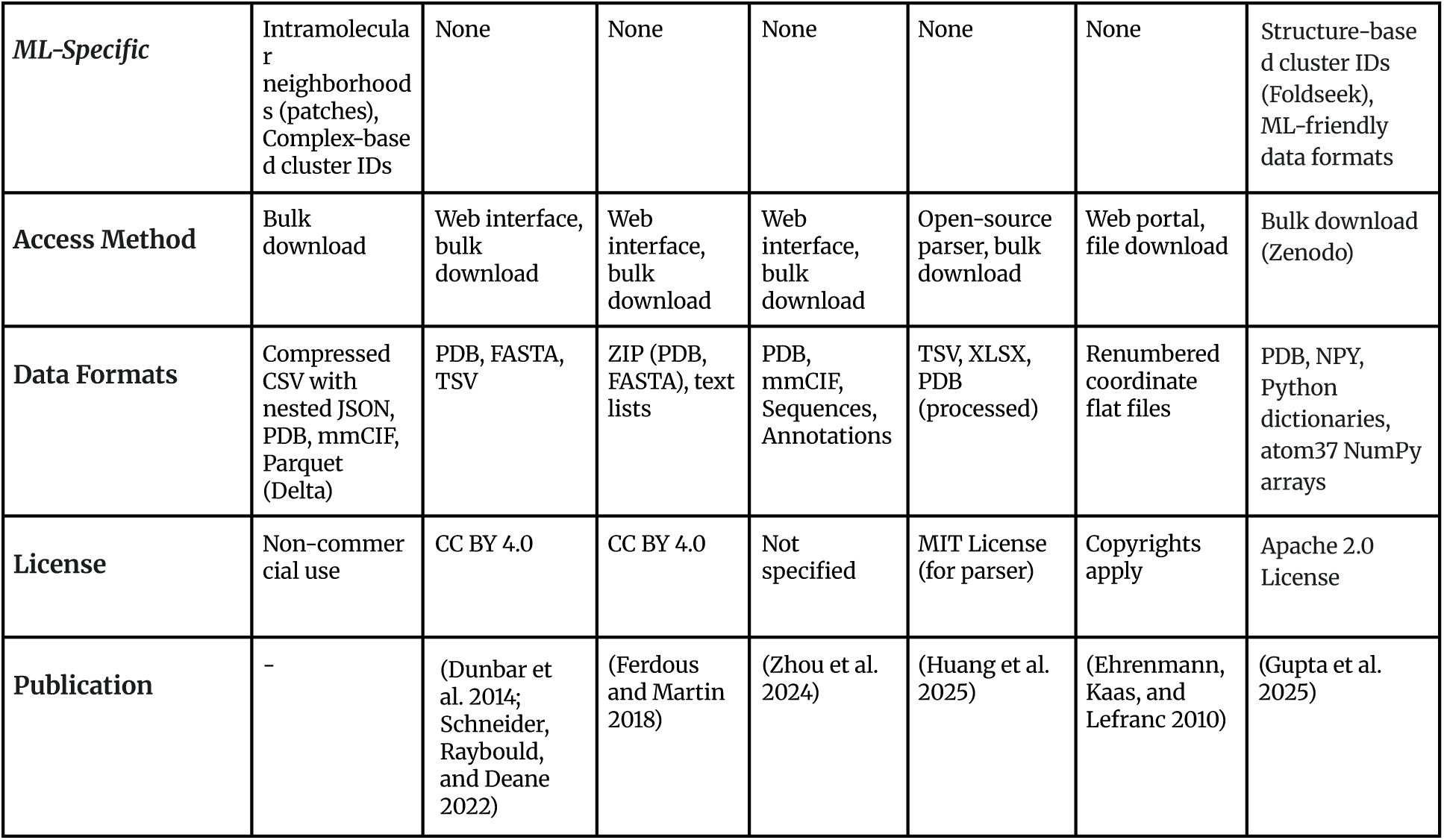
Feature comparison of major structural interaction databases. A detailed side-by-side evaluation of available databases (NAStructuralDB, SabDab (Dunbar et al. 2014), AbDb (Ferdous and Martin 2018), AACDB (Zhou et al. 2024), SAAINT-DB (Huang et al. 2025), IMGT/3Dstructure-DB (Ehrenmann, Kaas, and Lefranc 2010) and SNAC-DB (Gupta et al. 2025)). The comparison covers essential attributes for computational research, including data scope, curation, size, redundancy control, interface definition, annotation depth, and data formats, highlighting the unique positioning of each resource.

The database identifies antibody and Nanobody®-containing PDB files using the Rapid Immunoglobulin Overview Tool (RIOT (Dudzic et al. 2024)), which annotates amino acid sequences from RCSB SEQRES sections and aligns them to their corresponding structural residues. We extract eight distinct datasets summarized in Table 2 in the form of a deduplicated and full dataset, which can be divided between proteins, antibodies, Nanobodies® and their complexes. Within each subset, we divide the information into chain-level information, intra-molecular residue level and pairwise inter-molecular information (Figure 1). The underlying database is subject to regular updates. Consequently, the values presented herein reflect the state of the database at the time of writing.

**Figure 1.**
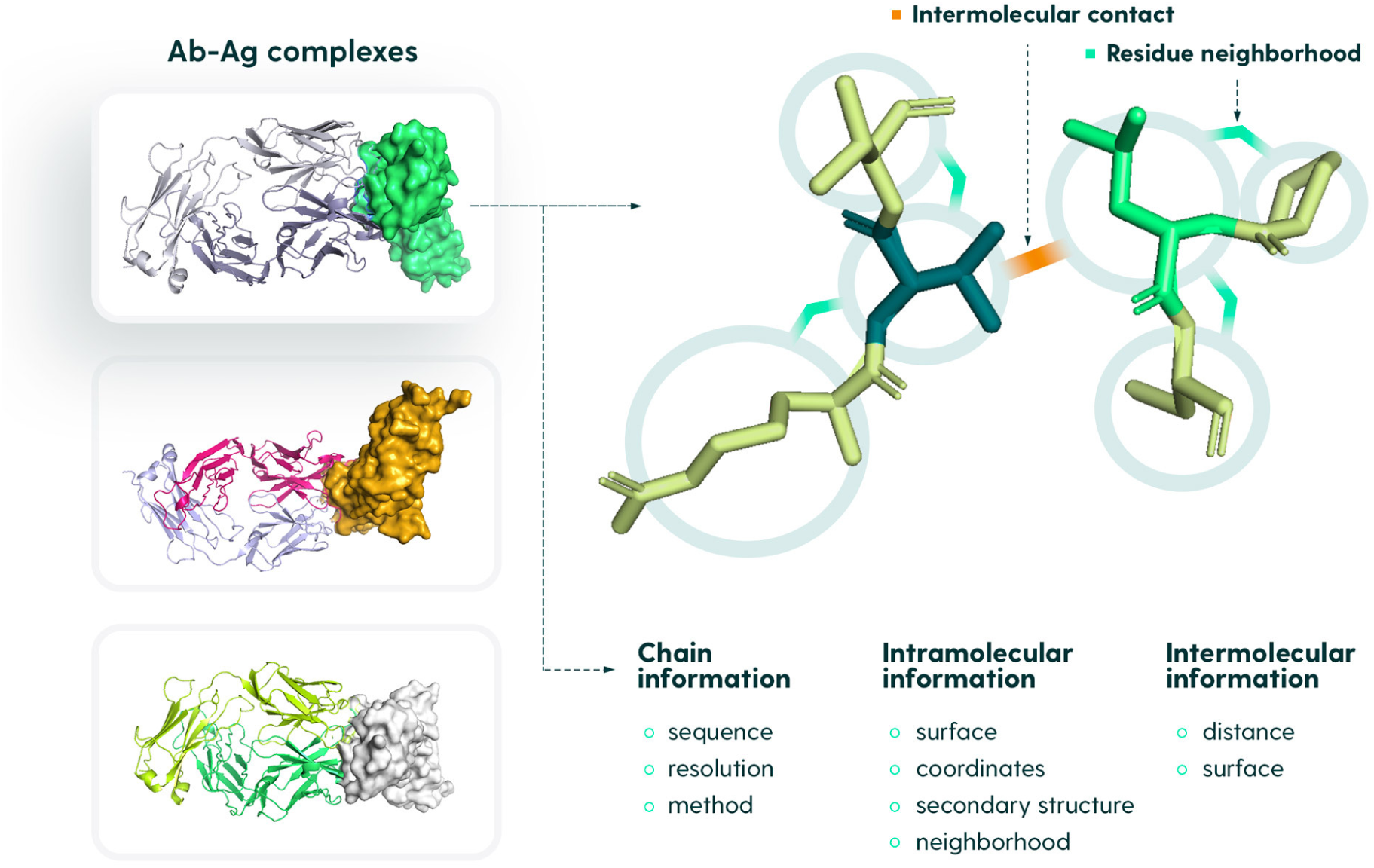
Database content. For each complex, we store the chain information, intramolecular information and intermolecular information. Chain information is used to perform filtering via sequence and experimental data quality. Intramolecular information is used to select paratope and epitope residues based on the contact distance to their binding partner, surface accessibility and secondary structure. Intermolecular contacts are pairs of residues between the paratope and epitope that are less than 10Å distance from one another heavy chain atoms. For subsets that do not contain complex information, the intermolecular information is not present.

**Table 2.** Datasets contained within NAStructuralDB. We divided NAStructuralDB into three separate molecule types: Nanobodies®, antibodies and other proteins. We cluster each of the datasets to offer a non-redundant and full set of either unbound chains or complexes. Note that a higher number of structures than the total number of PDBs reflects distinct chains/complexes within a single PDB file.

| Molecule type | Database | Number of structures - full dataset (October 2025) | Number of representative structures (October 2025) | Representative clustering procedure |
| --- | --- | --- | --- | --- |
| Nanobodies® | Unbound Nanobodies® | 4160 (from 2172 PDBs) | 884 (from 839 PDBs) | 90% sequence identity unique (cl_90_id) |
|  | Nanobody®-antigen complexes | 2768 (from 1469 PDBs) | 799 (from 752 PDBs) | 90% sequence identity of Nanobody®, 30% sequence identity of protein<br>unique ({cl_90_id, cl_30_id}) |
| Antibodies | Unbound antibodies (Paired heavy and light chains) | 13660 (from 7146 PDBs) | 3681 (from 3571 PDBs) | 90% sequence identity of heavy chain, 90% sequence identity of light chain<br>unique ({H:cl_90_id, L:cl_90_id}) |
|  | Antibody-antigen complexes | 8553 (from 4050 PDBs) | 2671 (from 2514 PDBs) | 90% sequence identity of heavy chain, 90% sequence identity of light chain, 30% sequence identity of protein<br>unique ({H:cl_90_id, L:cl_90_id, Ag:cl_30_id}) |
|  | Unbound heavy chains | 13815 (from 7189 PDBs) | 2613 (from 2559 PDBs) | 90% sequence identity of heavy chain<br>unique ({H:cl_90_id}) |
|  | Unbound light chains | 14412 (from 7391 PDBs) | 717 (from 707 PDBs) | 90% sequence identity of light chain<br>unique ({L:cl_90_id}) |
| Proteins | Unbound proteins | 895371 (from 230435 PDBs) | 32829 (from 27690 PDBs) | 30% sequence identity<br>unique<br>({cl_30_id}) |
|  | Holo proteins<br>(Hetero dimers) | 76599 (from 23954 PDBs) | 8379 (from 6737 PDBs) | 30% sequence identity of protein chain,<br>30% sequence identity of protein chain<br>unique<br>({P1:cl_30_id,<br>P2:cl_30_id}) |

Each dataset is available in the following formats:

● CSV / JSON - gzip compressed csv file with a row containing basic entry information and comprehensive json structure divided into basic, pairwise, protein1 and protein2 (if applicable) sections
● Parquet (delta) - data pipelines-enabled format with well-defined data structure
● PDB/mmCIF - structure derived from computed complex or chain in PDB and mmCIF format (available for deduplicated datasets).

*Chain level information*.

The chain level of abstraction in the data was designed to identify subsets of structures by offering ways to stratify by entire molecule sequence diversity and by quality of experimental data.

In the case of protein and Nanobody® unbound chain datasets, only single chain information is provided. In the case of antibodies without the antigen, information on both the heavy and light chain is provided. In the case of holo proteins and Nanobody®-antigen complexes, information on both interacting chains is given, whereas for antibodies the heavy, light and antigen information are present (Table 3).

**Table 3.** List of chain-level information entries. Each dataset contains dataset-type-specific information with fields related to available molecules inside the dataset. Below we present a generic description of available fields with the following possible molecule_types: *antigen, heavy, light, nanobody, chain1, chain2*.

| Field name | Field description |
| --- | --- |
| {molecule_typeX} | {molecule_typeX} chain id |
| {molecule_typeX}_seq | {molecule_typeX} chain sequence |
| {molecule_typeX}_organism_name | {molecule_typeX} organism name |
| {molecule_typeX}_taxonomy_id | {molecule_typeX} taxonomy id of organism |
| {molecule_typeX}_host_organism_name | {molecule_typeX} host organism name |
| {molecule_typeX}_host_taxonomy_id | {molecule_typeX} host taxonomy id of organism |
| cl_id | Compound cluster id (underscore concatenated heavy_cl_id, light_cl_id, antigen_cl_id) |
| l3_h3 | cdrl3, cdrh3 sequence joined by “_”. Data available if applicable |
| method | Experimental method |
| resolution | Experimental method resolution |
| last_update | Last update date |
| initial_release | Initial release date |
| deposition_date | Deposition date |

To allow for sequence redundancy removal, for each antibody/Nanobody® chain we offer the extracted sequence of CDR-H3 and CDR-L3. In addition, we provide complex-based redundancy removal by using the clustering results reference provided by RCSB.org at 90% sequence identity for antibody chains and 30% sequence identity for non-antibody chains. Each database entry is then associated with the underscore-concatenated cluster id. Deduplicated (non-redundant) datasets are constructed by selecting only a single member of the corresponding cluster_id sorted by a resolution quality and next by deposition date. Thus the deduplicated dataset contains the most recent top quality representatives of the cluster. By handling cluster_id chain-corresponding parts, sequences of different antibodies that target the same/similar antigen can be assigned to the same group in the train/validation/test splits, preventing information leakage. While our sequence-based clusters are a robust standard, for tasks requiring strict structural generalization we recommend an additional layer of structure-based clustering to ensure the most stringent train/test separation and prevent leakage from sequentially dissimilar but structurally related examples.

#### Intramolecular residue information

At the intramolecular information level, we retain data that should help identify contacting residues on the surface together with their neighborhoods (Table 3). Each residue is referred to by the chain id and the sequence positional id. Basic information such as residue type, residue ID within the structure and atomic coordinates are given. This information should facilitate navigating the molecule as well as recreating the entire PDB files, thus rendering our data format self-sufficient.

For antibody chains there are supplementary attributes, namely the IMGT id and the region (e.g. framework 1, CDR1). Both antibody chains are treated as a single molecule and thus are not separated. This facilitates treating the antibody as a single entity contacting the antigen chain as well as extracting information about the VH/VL interface.

Our data processing pipeline relies exclusively on the atomic coordinates provided within the ATOM records of the source Protein Data Bank (PDB) files. Consequently, all derived annotations, including intermolecular contact distances, interface definitions, surface accessibility calculations and secondary structures assignments, are based strictly on the atoms present in the original PDB entry. Users should be mindful that these calculations reflect the completeness of the source structure and may not encompass regions that were not resolved in the original experiment.

Pairing of the heavy and light chains has been motivated by earlier work from the Structural Antibody Database, where the heavy and light chain residue 104 (IMGT scheme) representing highly conserved cysteines - forming disulfide bonds - need to be less than 20 Å apart (Dunbar et al. 2014). Each combination of heavy and light chains is associated with the closest antigen chain. There is some ambiguity in the Protein Data Bank as to what antigen chain an antibody recognizes. Identification of the antigen chain closest to the antibody CDRs is a good approximation but not universally true as antigens can span multiple chains. Inferring such information from biological assemblies is also error prone - for instance 1ahw is a good example, where one biological assembly contains two copies of the same antibody. To select proper pairs of antibody antigens we require that at least one C-alpha or C-beta atom from the CDR region be less than 7.5 Å apart from antigen C-alpha or C-beta atoms.

To handle complex structural scenarios and ensure data integrity, our processing pipeline incorporates the following rules. For PDB entries that contain multi-chain antigens or multiple distinct Fabs, each antibody-antigen interaction is identified and stored as a separate entry in the database. For symmetric assemblies, our complex-based clustering protocol effectively handles deduplication; the full dataset provides all the interaction examples, while the representative dataset contains a single, canonical instance from each cluster, preventing artificial inflation of the data from crystallographic or biological symmetries.

Further information at this level includes surface, secondary structure and contact information. Table 8 contains references to the annotation tools used in the processing pipeline. Contact is given as both C-alpha distances or closest heavy atoms (Table 4). The contact field is capped at 10Å.

**Table 4.**
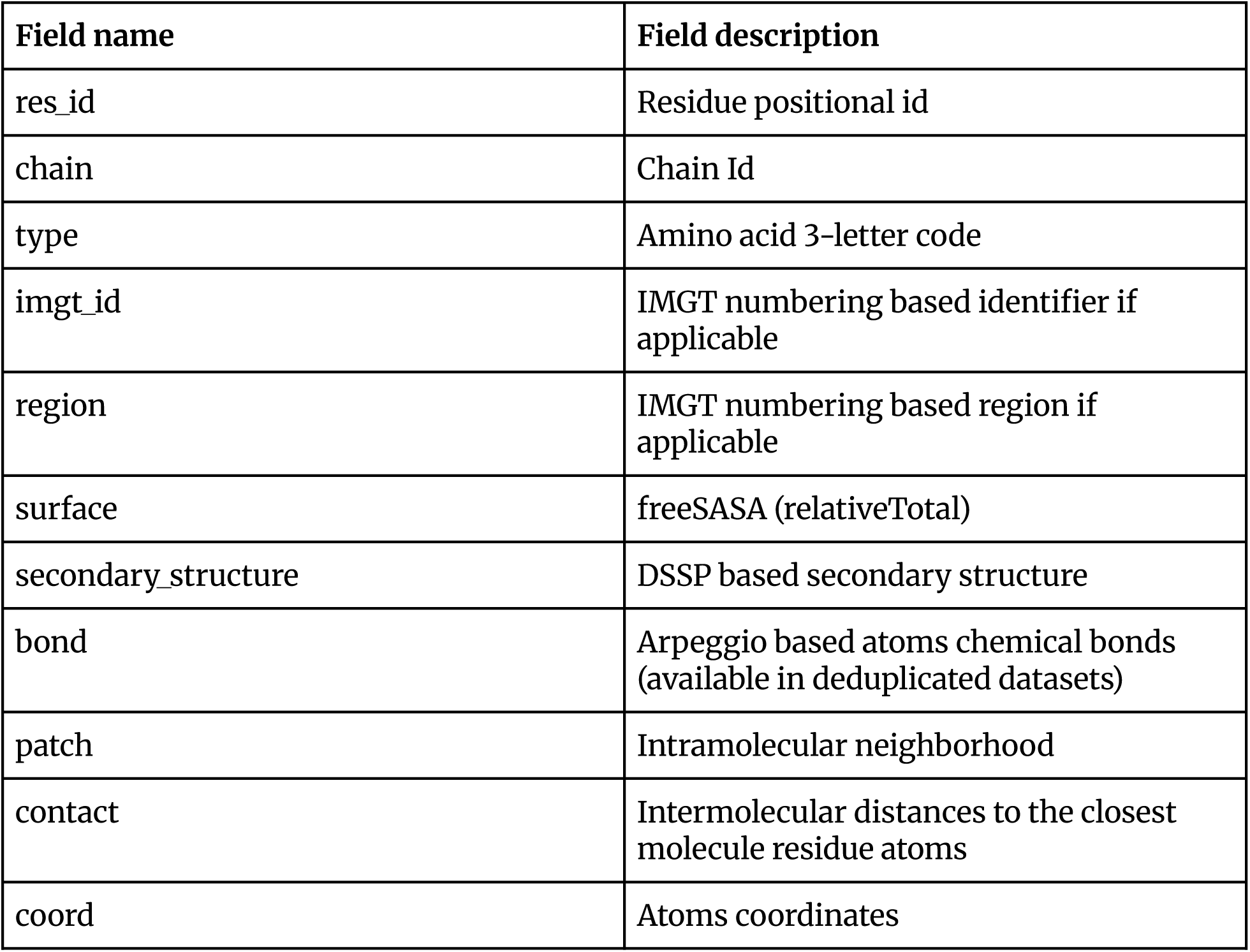
List of data fields for residue-level intramolecular information available inside the “protein1” and “protein2” (if applicable). Data structure is a key-value map with a key referred by underscore concatenated chain_id and a residue positional id eg. “H_101” and values with the fields.

| Field name | Field description |
| --- | --- |
| res_id | Residue positional id |
| chain | Chain Id |
| type | Amino acid 3-letter code |
| imgt_id | IMGT numbering based identifier if applicable |
| region | IMGT numbering based region if applicable |
| surface | freeSASA (relativeTotal) |
| secondary_structure | DSSP based secondary structure |
| bond | Arpeggio based atoms chemical bonds (available in deduplicated datasets) |
| patch | Intramolecular neighborhood |
| contact | Intermolecular distances to the closest molecule residue atoms |
| coord | Atoms coordinates |

Surface accessibility is an important metric when identifying paratope/epitope residues that are in contact and on the surface. It is counted as the % total relative surface area (tRSA), also known as relative solvent accessibility. It is a measure of a protein residue’s exposure to solvent of all atoms in the amino acid and calculated by freeSASA (Mitternacht 2016), in the absence of the binding partner (but for antibody heavy and light chains together). It is calculated using equation 1, with *maxASA* reference value derived from Ala-X-Ala tripeptides in an extended conformation:

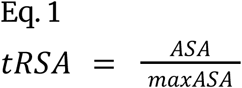

While it can be counterintuitive, the occurrence of total relative solvent accessibility (tRSA) values exceeding 100% when using FreeSASA can be attributed to the reference values used for normalization. FreeSASA calculates RSA by comparing the observed solvent-accessible surface area (SASA) of a residue to a reference maximum SASA value. These reference values are derived from Ala-X-Ala tripeptides in an extended conformation. However, since these configurations may not represent the absolute maximum exposure possible for each residue, and due to variations in bond lengths and angles, the observed SASA can sometimes surpass the reference values, resulting in RSA values greater than 100%. Computed values are kept unchanged in the database.

At the intramolecular level we are also facilitating extraction of the molecular neighborhoods (Table 6). Such information enriches the data content of the individual interacting residues, by prediction of patches (Tubiana, Schneidman-Duhovny, and Wolfson 2022). Patch information for each residue includes the residues that are within 8Å of it by a heavy atoms distance. It also includes the distances as described in Table 5 and surface accessibility information to allow for introduction of different filters for patch inclusion.

**Table 5.** List of distance structure fields.

| Field name | Field description |
| --- | --- |
| sidechain_and_ca_atoms | Closest distance between residues sidechain and CA atoms |
| ca_atoms | Closest distance between residues CA atoms |
| heavy_atoms | Closest distance between residues heavy atoms (all atoms without hydrogens) |

**Table 6.** List of value fields for patch of intramolecular information. Each patch entry is referenced by chain id and the sequence positional id.

| Field name | Field description |
| --- | --- |
| distance | Structure describing distance to this residue |
| imgt_id | IMGT numbering based identifier if applicable |
| surface | freeSASA (relativeTotal) |

#### Intermolecular residue information

We store intermolecular information to facilitate retrievals for contact predictions, such as antibody-antigen docking. Each entry in the pairwise information contains data on the identifiers of residues from the interacting partners and their amino acid types (Table 7). Most importantly, we store the closest side chain and C-alpha distances, closest C-alpha distances and closest heavy atoms distances in Ångstroms between the two amino acids. For convenience we also store the surface accessibility area. We only keep the pairs where the closest distance is less than 10Å.

**Table 7.**
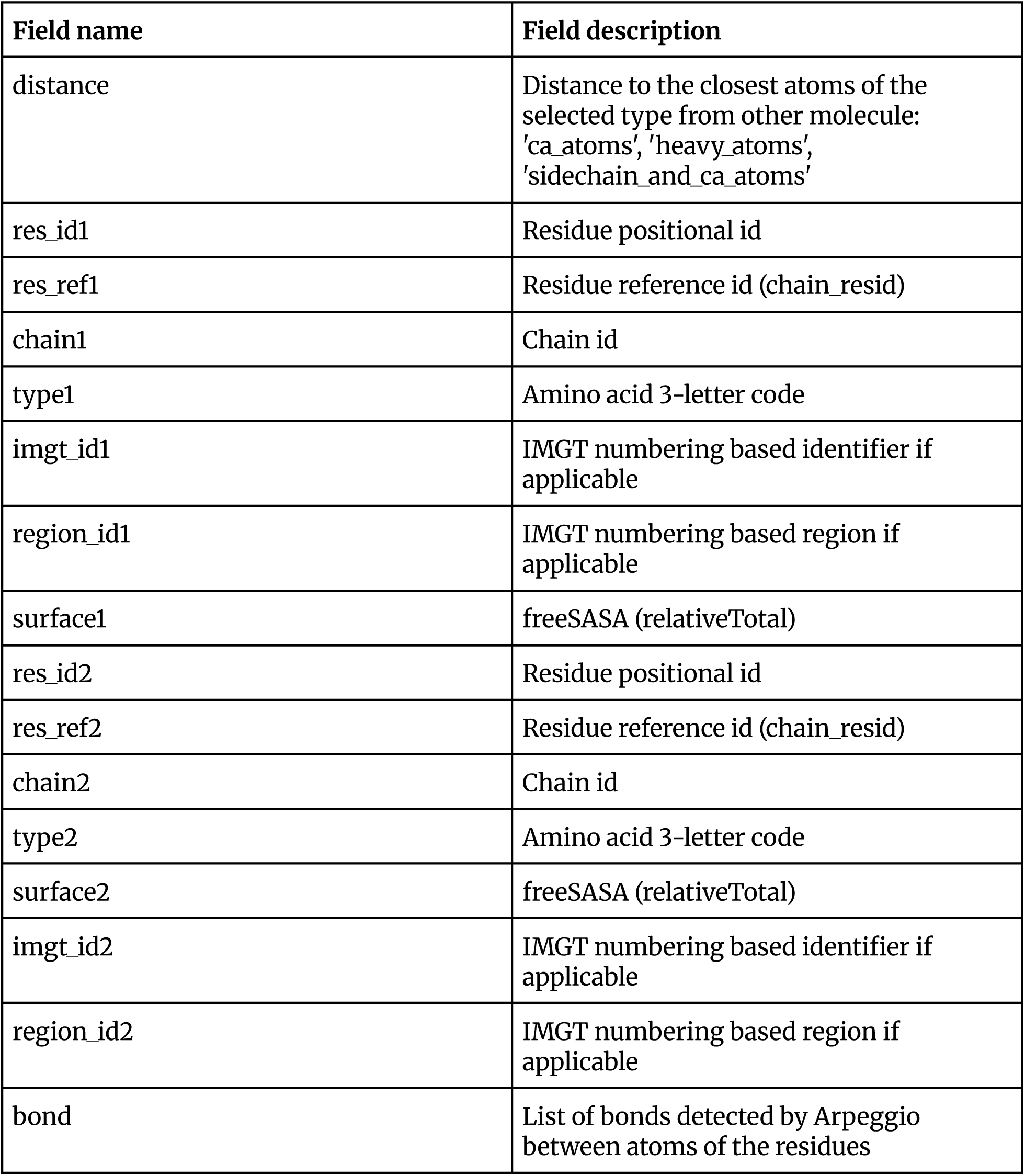
List of data fields for intermolecular information. All of the intermolecular distances are holding references for each derived bioassembly in the form of bio_assembly_id - intermolecular information mapping with −1 as a default model definition.

| Field name | Field description |
| --- | --- |
| distance | Distance to the closest atoms of the selected type from other molecule: 'ca_atoms', 'heavy_atoms', 'sidechain_and_ca_atoms' |
| res_id1 | Residue positional id |
| res_ref1 | Residue reference id (chain_resid) |
| chain1 | Chain id |
| type1 | Amino acid 3-letter code |
| imgt_id1 | IMGT numbering based identifier if applicable |
| region_id1 | IMGT numbering based region if applicable |
| surface1 | freeSASA (relativeTotal) |
| res_id2 | Residue positional id |
| res_ref2 | Residue reference id (chain_resid) |
| chain2 | Chain id |
| type2 | Amino acid 3-letter code |
| surface2 | freeSASA (relativeTotal) |
| imgt_id2 | IMGT numbering based identifier if applicable |
| region_id2 | IMGT numbering based region if applicable |
| bond | List of bonds detected by Arpeggio between atoms of the residues |

#### Quantitative Metrics for Interface and Surface Composition Analysis

To quantitatively assess the compositional biases of amino acid residues at protein surfaces and binding interfaces, we defined a series of metrics. These calculations allow for a systematic comparison of residue propensities between different interaction types, such as general protein-protein interactions (PPIs) and antibody-antigen (Ab-Ag) complexes.

#### Single Residue Propensity Metrics

For the purpose of single residue analysis, let s_i_ be the number of residues on the surface of amino acid type *i*, aa_i_ the total number of residues of the amino acid type *i* (7.5% relative side chain exposure), and c_i_ the number of residues in contact of amino acid type *i* (4.0Å heavy atom distance). The following metrics were calculated:

- ● Surface Rate(AA) - The extent to which a residue is found on the surface. It is calculated as the ratio of the occurrence on the surface versus the total number of residues:

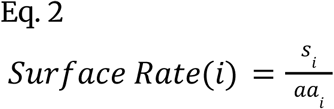

- ● Contact Rate on Surface(AA) - The extent to which a residue is in contact, given that it is on the surface:

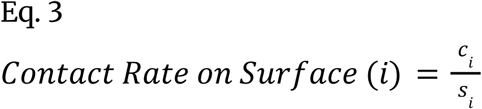

- ● Contact Frequency(AA) - The propensity of a certain residue type to form contacts. Calculated as the frequency of a residue type across all residues in contact:

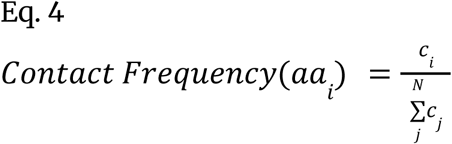

- ● Normalized Contact Frequency(AA) - The log_2_ proportion of a residue type in contact relative to its overall frequency across all residues. It is calculated with equation 5 using Frequency defined in equation 6:

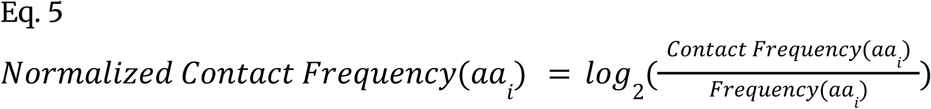

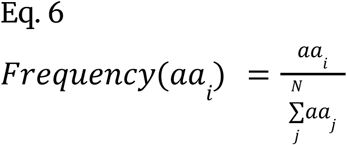

#### Residue Pair Contact Propensity Metrics

To analyze the preferences for specific residue-residue pairings at interfaces, we first calculated the following basic statistics:

● c_ij_: the number of contacts between residues AA1 and AA2
● c_i_ : the total number of contacts involving residue type AA1,
● c_j_: the total number of contacts involving residue type AA2 Based on these statistics, we defined the following metrics:
● **Contact Frequency(AA1_AA2)** - represents how often residue type AA1 is observed in contact with residue type AA2 across all residues participating in interfaces:

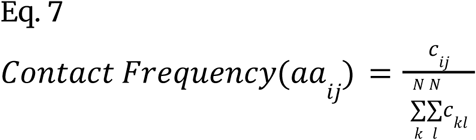

- ● **Normalized Contact Frequency(AA1_AA2)** - adjusts the contact frequency between AA1 and AA2 by accounting for the overall frequencies of AA1 and AA2 in the interface. This normalization highlights residue pairs that are observed in contact more (or less) often than expected by chance, based on the independent frequencies of each residue type:

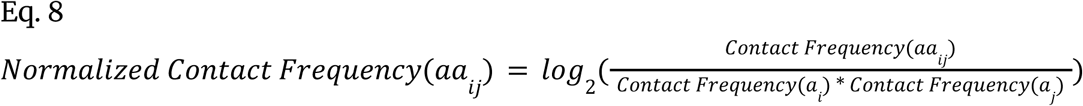

### Database access

The database is freely available for non-commercial organizations for non-commercial purposes at https://naturalantibody.com/na-structural/.

## RESULTS

### Antibody antigen contact analysis and reference

For further analysis we took a February 2025 release of an antibody-antigen complex database containing 1172 entries from 1136 PDBs filtered to high quality (<3Å resolution) X-ray diffraction structures and performed a survey of antibody-antigen contacts and contrasted them to protein-protein interfaces, which has been attempted several times before albeit with smaller datasets (Krawczyk et al. 2013; Kunik and Ofran 2013; Mitchell and Colwell 2018b, 2018a; Reis et al. 2022). From the protein-protein dataset we extracted the hetero-dimers with deduplication using 30% sequence identity and processed them using the same pipeline as for antibody-antigen complexes resulting in 5158 unique complexes from 4453 PDBs (we do not consider hetero-dimers only, hence the greater number of complexes than PDBs).

### How far away are residues from one another in inter- and intramolecular interactions?

One of the chief considerations when performing contact prediction is to define the atomic distance cutoff. Some of the commonly employed definitions are 1) C-alpha distance 2) closest heavy atom (including backbone) 3) closest heavy side chain atom (with special treatment for glycine). Depending on the definition used, the cutoff also needs to be adjusted - for instance, the distance cutoff between C-alpha atoms would on average be larger than the distance cutoff between the side chain atoms. Nevertheless, many different cutoffs are used throughout the literature, ranging from 1.5Å to 8Å within a single definition.

To provide a point of reference, we calculated the proportions of contacting residues for a range of cutoffs using our definition of distances, which are 1) the closest heavy atoms distance and 2) the closest C-alpha atoms distance. The top panels in Figure 2 show the average number of contacts for antibody/antigen and protein dimers for a range of cutoffs between 2.5Å and 5.0Å for heavy atoms and 4.0Å to 12.0Å for C-alpha atoms. Both antibody/antigen as well as protein dimers appear to have a similar number of contacts until a cutoff of 3.5Å for heavy atoms distance, at which stage protein dimers start having more contacts. In case of C-alpha distances, a similar discrepancy happens at a distance of approximately 5Å. The heavy atoms distance graphs appear to have a sigmoidal shape with the steepest interval between approximately 3Å and 4Å, while for C-alpha atoms the distance graph is more linear.

**Figure 2.**
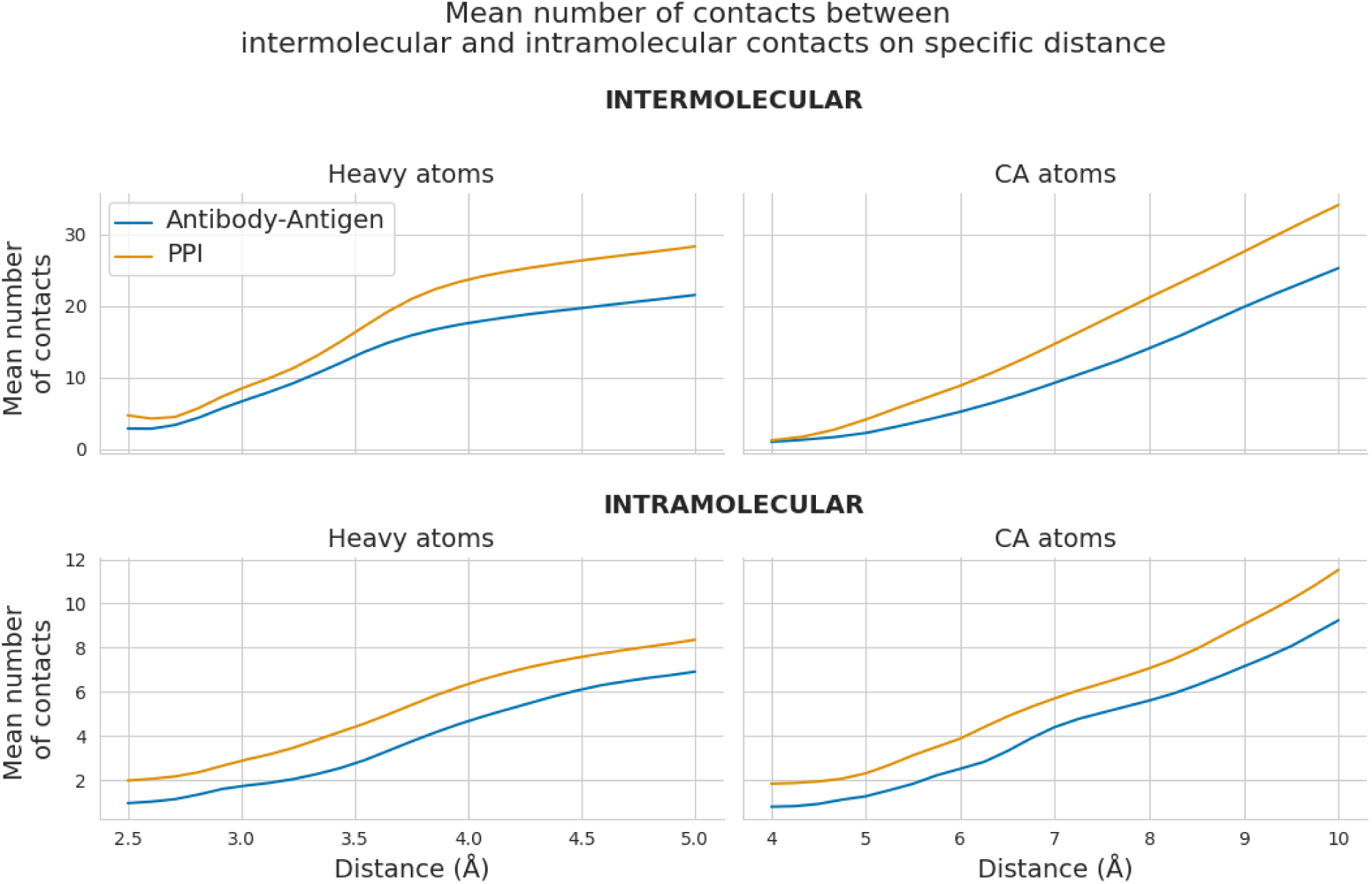
**Distribution of contact distances of intermolecular and intramolecular contacts. Top Left**: intermolecular surface contacts, with distance computed using the closest heavy atom. **Top Right**: intermolecular surface contacts, with distance computed using the closest C-alpha. **Bottom Left**: intramolecular surface contacts, distance computed using the closest heavy atom. **Bottom Right**: intramolecular surface contacts, distance computed using the closest C-alpha.

To compare with the intermolecular contacts, we also investigated distances between intramolecular contacts (Figure 2 bottom panels). Both heavy-atom distance and C-alpha distance based metrics give similar shapes, however with proteins being more compact with the higher number of contacts for the same distance cutoffs as antibodies. Heavy-atom distance recapitulates the sigmoidal shape whereas C-alpha distance is more linear for both.

To compare with the total number of contacts, we also plotted the distance at which there are no contacts at all (Figure 3). For both antibody/antigen as well as protein dimers there appears to be an inflection point at around 2.5Å for heavy atoms and 4Å for C-alpha distances. For distances lower than those inflection points there are no contacts and beyond there is a rapid accumulation of those. These can certainly be explained by atomic radii and delineating it like we have done here gives a reference for what is a lower bound of difference to be considered a contact altogether.

**Figure 3.**
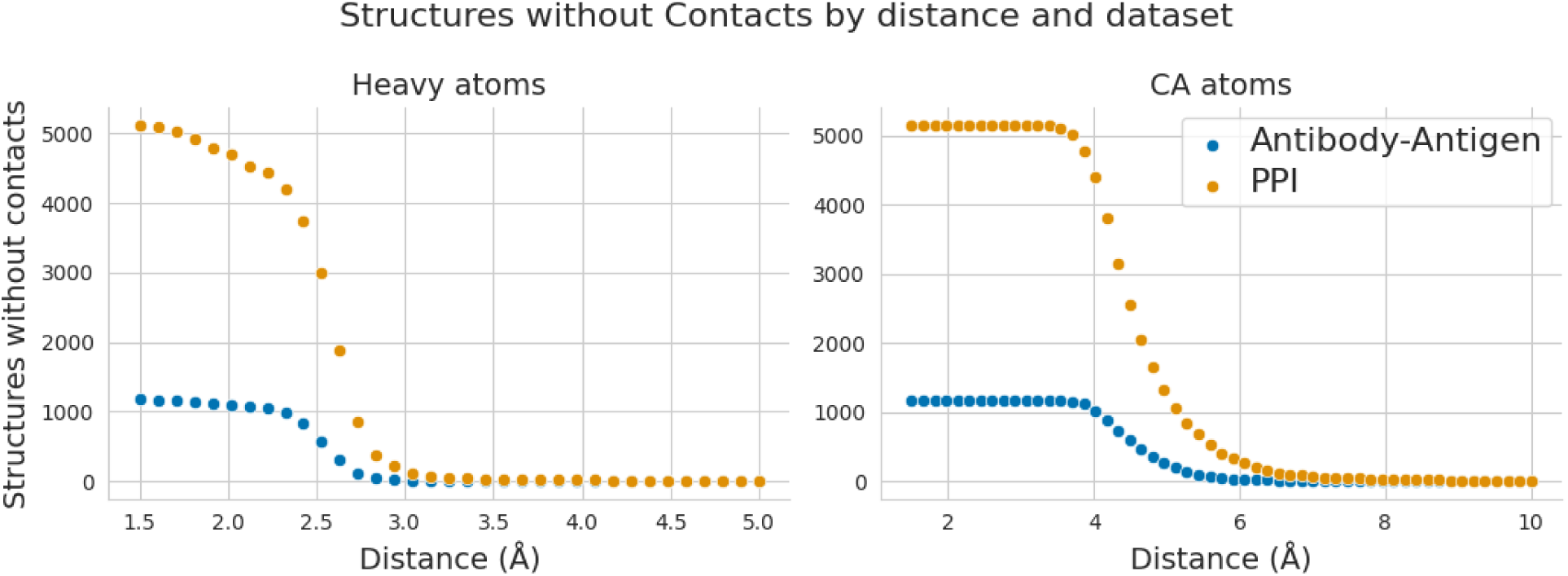
Number of complexes without any contacts detected at a specific threshold for antibody-antigen complexes and protein protein complexes (PPI). **Left**: heavy atoms as a distance metric. **Right**: C-alpha atoms as a distance metric.

### Total relative surface area (tRSA) thresholds comparison

Proteins chiefly interact through residues on their surfaces, which requires an understanding of how much of a protein chain is exposed on average. We plotted the number of residues in a protein (for both the PPI and Antibody-Antigen datasets) versus the number of surface exposed residues in Figure 4. Mean RSA, which represents the average extent to which residues are exposed to solvent, decreases as protein size increases. This trend suggests that larger proteins have a greater proportion of buried residues, leading to lower overall solvent exposure. The dataset comparisons indicate that while both follow a similar pattern, the Antibody/Antigen dataset has a denser distribution at lower protein residue counts.

**Figure 4.**
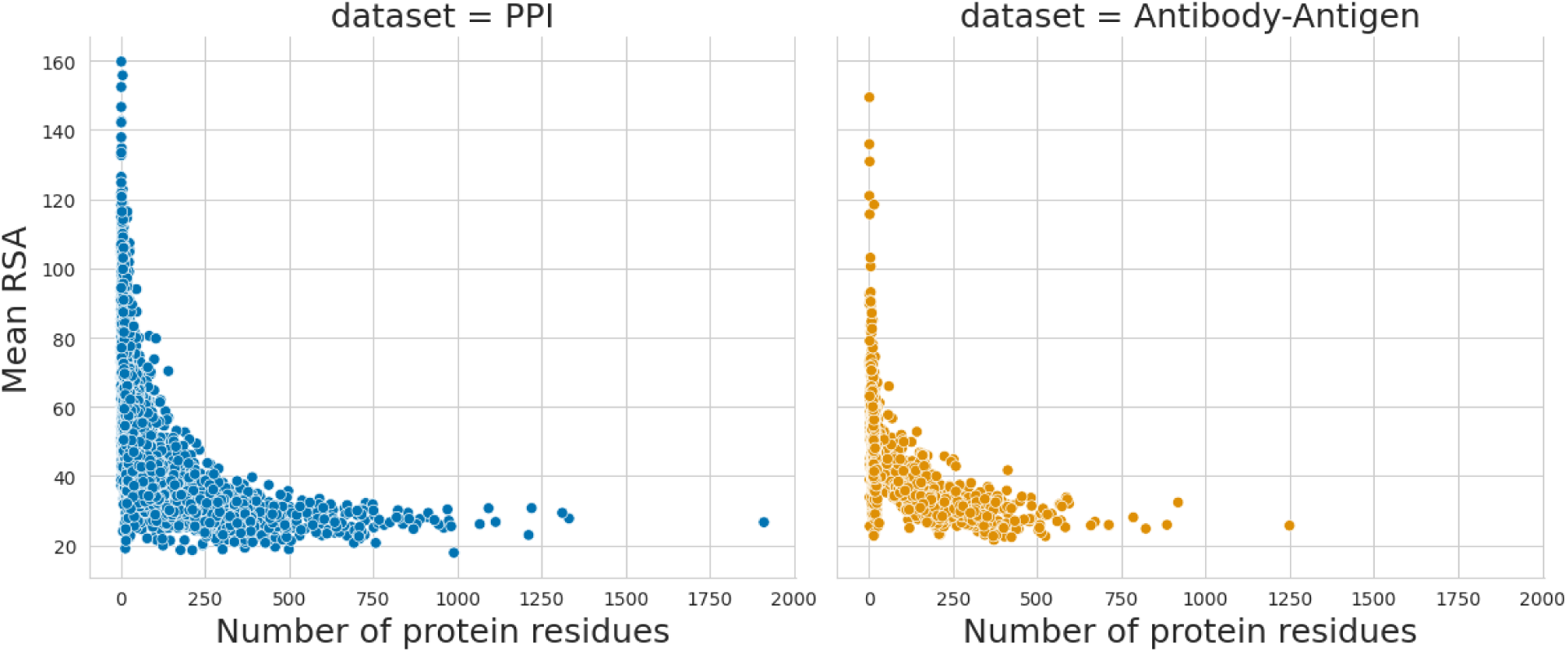
Mean relative surface accessibility of the protein residues. The surface exposed residues were taken as those with more than 7.5% of their surface exposed.

The number of buried residues shows the same strong linear correlation with protein size across different thresholds of solvent accessibility. As the threshold defining “buried” residues increases (from 7.5% to 30%) as shown on Figure 5, the absolute number of buried residues grows, but the overall trend remains consistent. This suggests that the fraction of residues sequestered from the solvent is proportional to protein length, irrespective of the specific threshold used. The PPI dataset exhibits greater variability, possibly due to the structural diversity of proteins involved in protein-protein interactions, whereas the Antibody/Antigen dataset follows a more constrained distribution.

**Figure 5.**
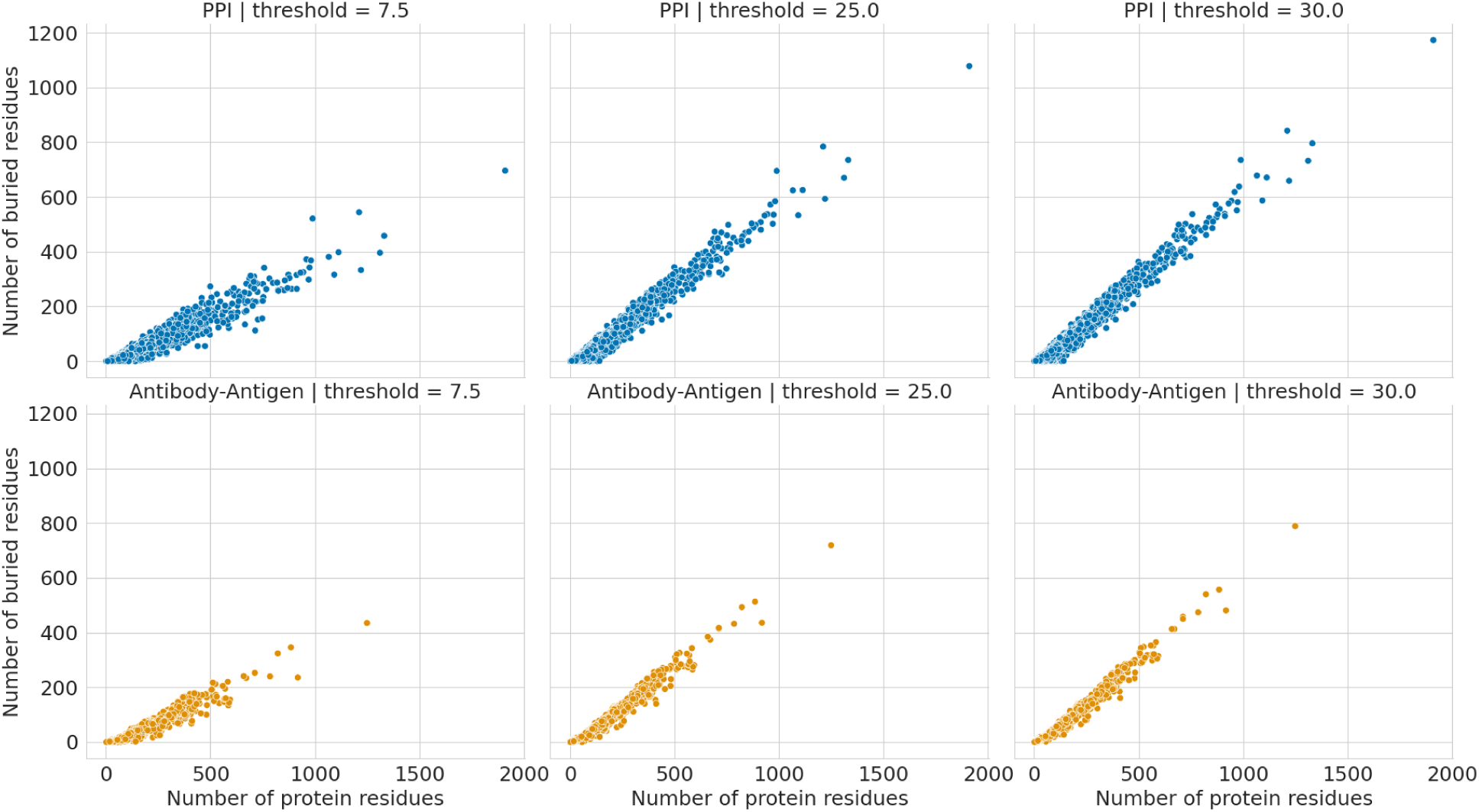
Comparison of number of buried residues for specific threshold values to the number of residues in protein used for relative surface accessibility calculations.

These findings align with fundamental principles of protein folding and stability, where larger proteins tend to have a compact core with lower solvent accessibility, ensuring structural integrity. The observed dataset differences may reflect distinct structural constraints, such as the necessity for antibodies to maintain accessible binding sites while still exhibiting some degree of burial for stability.

To further analyze structural dataset patterns, a threshold of 7.5% relative solvent accessibility was selected to define buried residues. This threshold provides a biologically relevant, albeit arbitrary, criterion for distinguishing between surface-exposed and buried regions within a protein. A lower threshold ensures that only residues with minimal exposure contribute to the buried fraction, making the analysis more sensitive to core structural features. Additionally, the strong linear correlation observed at this threshold suggests that it effectively captures the relationship between protein size and buried residues while maintaining consistency across different datasets.

### Total interface area of antibody/antigen complexes versus non-antibody proteins

The interface buried surface area is calculated as the sum of the unbound surface area of residues in contact - we investigated how the buried surface area would change depending on the number of residues included in such a calculation. As shown in Figure 6, the largest increase in surface area occurs between distance thresholds 3Å and 4Å, capturing the primary, biologically relevant interactions. There is a smaller increase from 4Å to 5Å suggesting the inclusion of more peripheral contacts.

**Figure 6.**
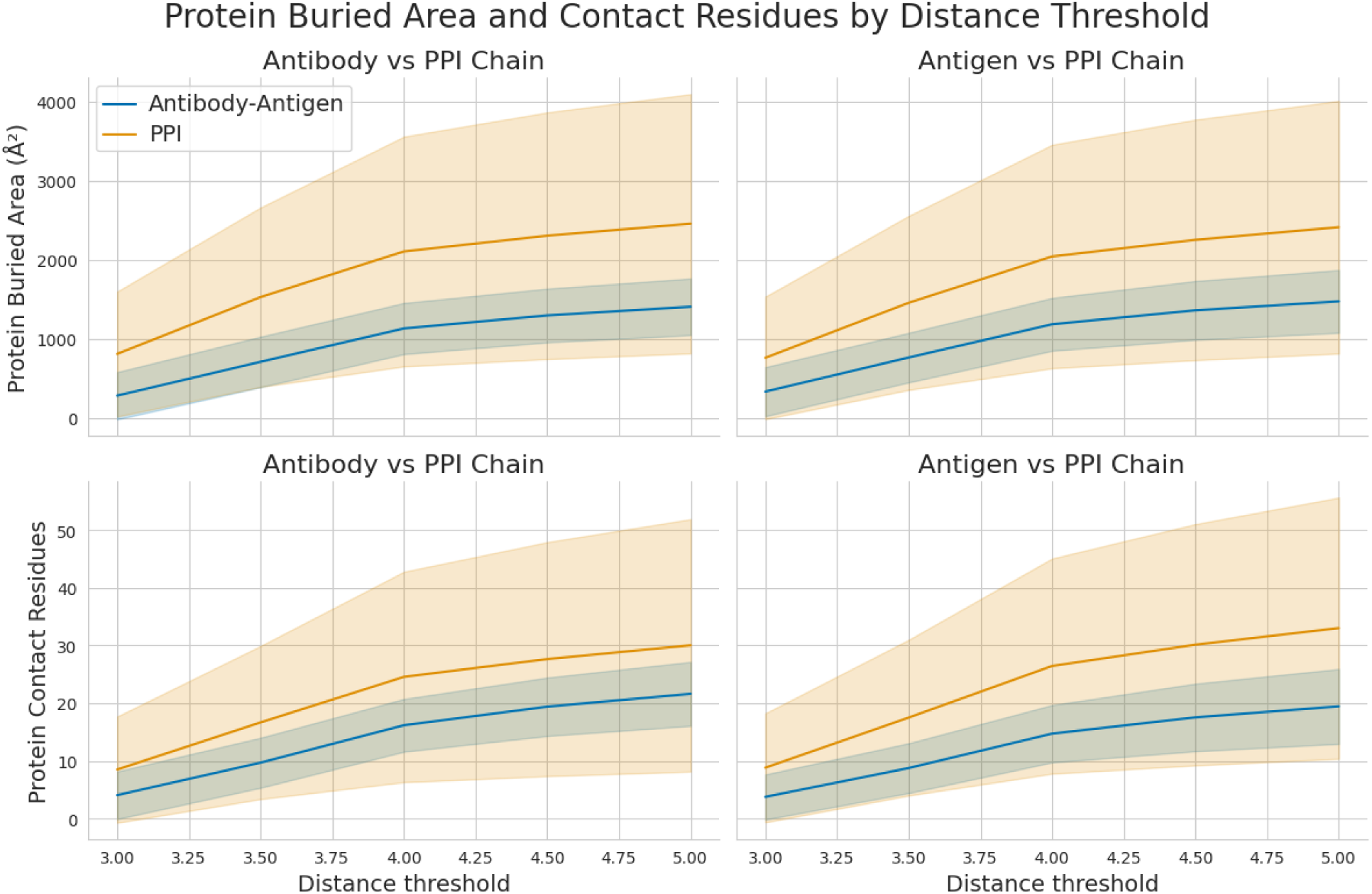
Comparison of protein interface area and contact residues by distance threshold. The figure compares antibody-antigen and protein-protein interaction (PPI) datasets. The top panels show the mean interface buried surface area (Å²), while the bottom panels show the mean number of contact residues, both plotted against increasing intermolecular distance thresholds. The shaded regions represent the variability within each dataset calculated by standard deviation. The largest increase in both metrics occurs between 3Å and 4Å, capturing primary biological interactions. The PPI dataset consistently shows greater variability, suggesting more diverse interaction modes compared to the more stable and constrained interaction profiles observed in antibody-antigen complexes.

It is also apparent that the PPI dataset captures a wider range of interactions, demonstrating greater variability in surface area and contact residues, as indicated by broader error bands calculated as standard deviation between complexes. This variability points to the more flexible and diverse interaction modes characteristic of PPI chains. On the other hand, antigen chains in the antibody-antigen dataset show more stable interaction profiles, with less fluctuation in these metrics. This could either point to a difference between protein surfaces versus interactions sites or indicate a smaller variability of the antigen-antibody dataset with respect to the larger PPI dataset.

### Differences between residue frequencies on interfaces vs surfaces

Certain residues would have a physicochemical preference to be either buried or on the surface, with the exposed ones being more likely to engage in interactions. To reveal such preferences, we calculated the frequency differences of residues that are buried versus those on the surface. Based on the metrics defined in Equations 2–6 of the Methods section, we calculated amino acid frequencies to analyze their distribution on protein surfaces versus their involvement in contact interfaces. The results, presented in Table 9, highlight the distinct compositional preferences that differentiate general protein-protein (PPI) from antibody-antigen interactions.

**Table 8.** List of external tools used for annotations in the processing pipeline.

| Tool | Version | Usage | URL | Publication |
| --- | --- | --- | --- | --- |
| DSSP | 4.0.4 | Secondary structure assignment using Kabsch-Sander standardized assignment algorithm | https://github.com/PDB-REDO/dssp | (Kabsch and Sander 1983; Hekkelman et al. 2025) |
| FreeSASA | 2.2.1 | Surface area calculation using Lee & Richards algorithm with Shrake & Rupley implementation. For the surface area annotation relativeTotal is being used, which is defined as the fraction of residue area related to Ala-X-Ala reference area. | https://github.com/freesasa/freesasa-python | (Mitternacht 2016) |
| PDBE Arpeggio | 1.4.4 (git) | Interatomic contacts calculation based on the rules defined in CREDO | https://github.com/PDBEurope/arpeggio | (Jubb et al. 2017) |
| RIOT | 4.0.2 | Amino acid immunoglobulin sequences annotation using an open germline database | https://github.com/NaturalAntibody/riot_na | (Dudzic et al. 2024) |

**Table 9.**
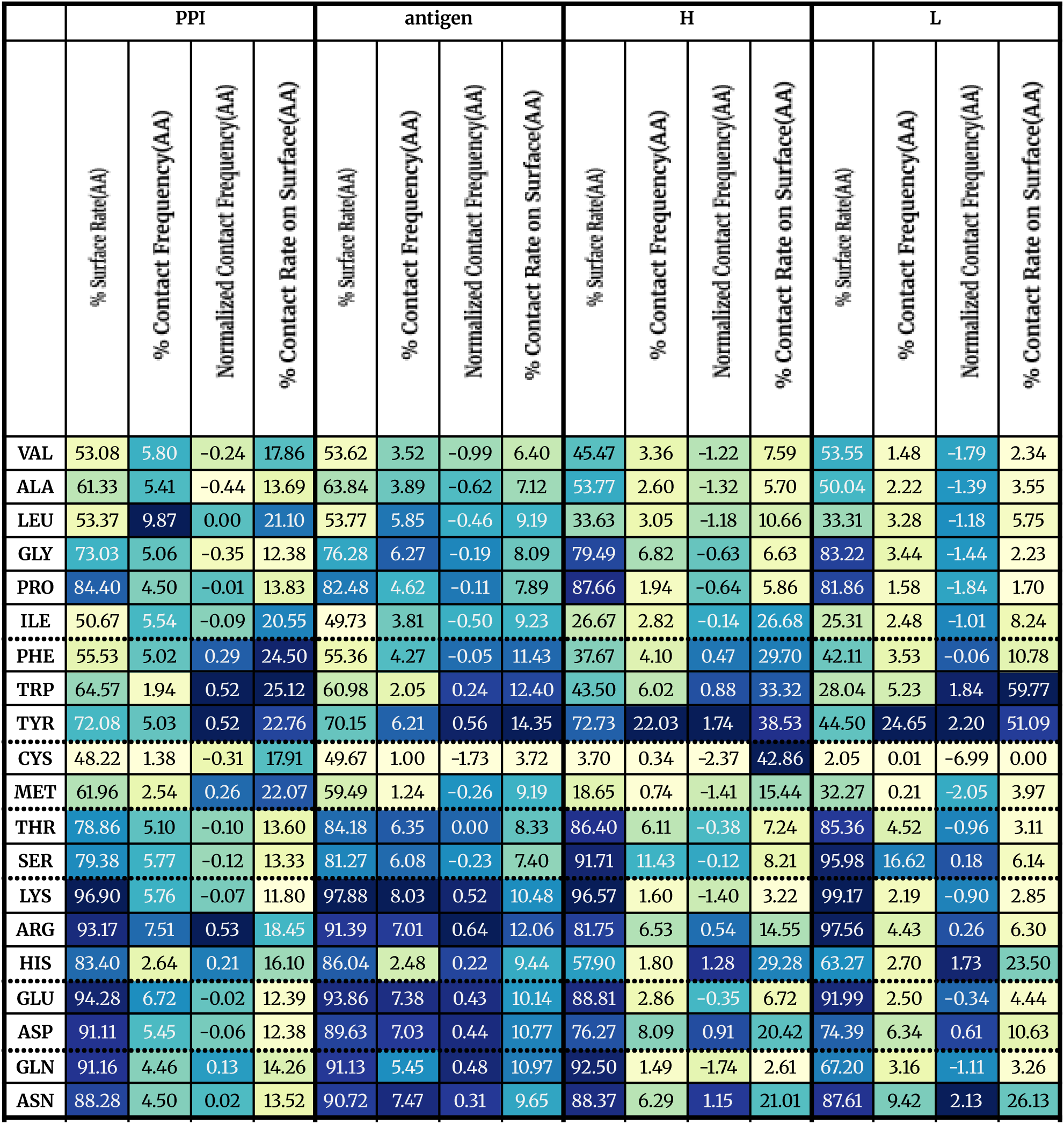
*Residue surface exposure and interface involvement in protein-protein (PPI) and antibody-antigen (Ab-Ag) complexes.* For each amino acid type across **PPI**, **antigen**, and antibody **heavy (H)** and **light (L)** chains, four metrics are shown: **surface rate (%), contact frequency (%), normalized contact frequency**, and **contact rate on surface (%)**. The normalized contact frequency indicates whether a residue is enriched (positive values) or depleted (negative values) at interfaces relative to its overall frequency. The data highlight consistent enrichment of aromatic (TYR, TRP) and polar (ASN) residues at interfaces and the depletion of hydrophobic (LEU, VAL, ILE) and cysteine (CYS) residues, in agreement with established trends in protein interface composition

The distribution of amino acid residues at protein-protein (PPI) and antibody-antigen (Ab-Ag) interfaces reveals conserved patterns that support the stability and specificity of molecular recognition. As shown in Table 9 and highlighted in Figure 7, aromatic residues, particularly tyrosine (TYR) and tryptophan (TRP), are highly enriched at interfaces. TYR displays consistently high normalized contact frequencies in both PPI (0.52) and antigen (0.56) datasets, with even stronger enrichment in the antibody heavy (1.74) and light (2.20) chains. Similarly, TRP shows significant contributions, especially within antibody interfaces. These observations confirm previous findings highlighting the dominant role of aromatic residues in stabilizing protein interfaces through π-π stacking, hydrogen bonding, and hydrophobic interactions (Mohamed, Degac, and Helms 2015; Chakrabarti and Janin 2002). In antibodies, the prevalence of TYR and TRP is particularly notable, reflecting their critical role in antigen recognition and binding versatility (Kunik and Ofran 2013; Reis et al. 2022).

**Figure 7.**
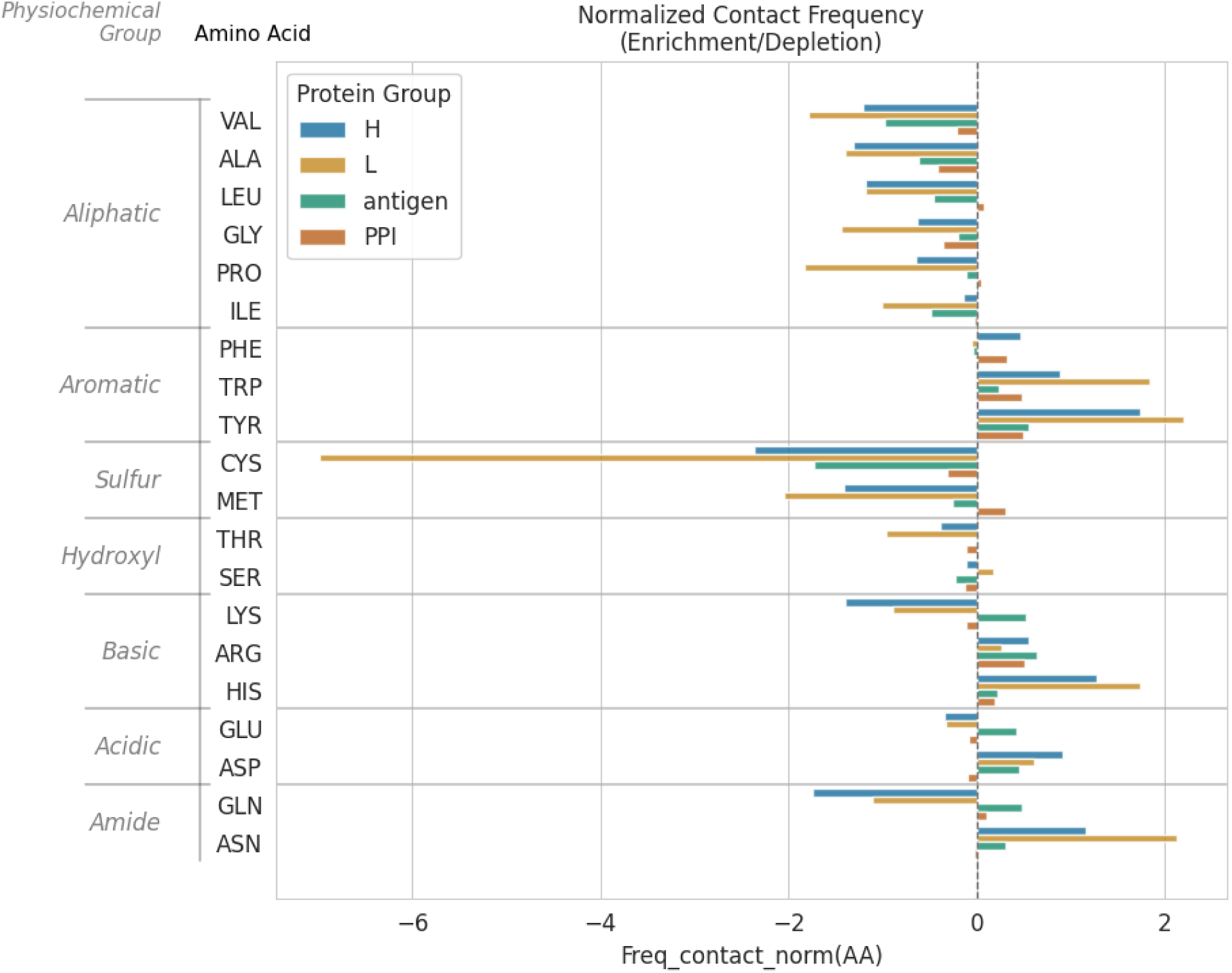
*Comparison of amino acid contact preferences across antibody-antigen and general protein-protein interfaces.* The normalized contact frequency (log ratio) indicates the enrichment (positive values) or depletion (negative values) of each amino acid relative to its expected abundance. The analysis reveals distinct compositional biases for antibody heavy (H) and light (L) chains and their corresponding antigens compared to a reference protein-protein interaction (PPI) dataset. Notably, aromatic residues, particularly Tyrosine (TYR) and Tryptophan (TRP), are enriched at the antibody-antigen interface, while Cysteine (CYS) is significantly depleted in the antibody-antigen chain contacts.

In contrast, cysteine (CYS) is markedly underrepresented across all interface types, as indicated by its consistently negative or near-zero normalized contact frequencies (−1.73 in antigen, −2.37 in heavy chains). This supports prior studies demonstrating that CYS primarily serves structural roles, such as forming disulfide bonds, rather than participating directly in dynamic protein-protein interactions (Hogg 2003).

Charged residues, notably arginine (ARG) and lysine (LYS), also show distinct patterns. ARG exhibits positive normalized contact frequencies across PPIs (0.53) and antigens (0.64), contributing to the electrostatic stabilization of complexes. While LYS is highly surface-exposed, its normalized contact frequency is variable, with reduced contributions in antibody interfaces (−1.40 in heavy chains), suggesting it may play a greater role in general surface solubility rather than in direct interface contacts within immune complexes.

Hydrophobic residues, including leucine (LEU), valine (VAL), and isoleucine (ILE), are generally depleted at interfaces. Their normalized contact frequencies are negative or near zero across all datasets (e.g., LEU 0.00 in PPI, −0.46 in antigen, −1.18 in heavy chains), confirming that these residues tend to be buried in the protein core, contributing to structural integrity rather than solvent-exposed interaction sites (Jones and Thornton 1996).

Polar residues such as asparagine (ASN) and threonine (THR) show more nuanced behavior. ASN, in particular, is enriched at antigen (0.31) and antibody interfaces (1.15 in heavy chains, 2.13 in light chains), reflecting its role in promoting hydrogen bonding and structural adaptability in antigen recognition. In contrast, THR remains relatively neutral across interfaces, with minimal contribution to binding, suggesting a supportive but non-dominant role.

These observations, supported by the data in Table 9 and highlighted in Figure 7, illustrate that aromatic and polar residues are fundamental to the formation and specificity of protein interfaces, particularly within adaptive immune responses, while hydrophobic residues remain predominantly buried and structurally focused. Charged residues further stabilize interfaces, particularly in general PPIs. Understanding these residue preferences is essential for advancing protein engineering, therapeutic antibody design, and interface prediction models, as they reflect the balance of forces driving molecular recognition and binding efficiency.

### Residue Pair Contact Analysis in Antibody-Antigen Interfaces Compared to PPIs

The analysis of residue-residue contacts was performed using a dual approach: calculating simple contact frequencies to identify the most common pairings, and a normalized contact frequency (log-odds score) to reveal specific interaction propensities by correcting for background amino acid abundance. While this normalization is crucial for uncovering true chemical preferences, the methodology is sensitive to statistical artifacts, particularly for rare amino acids. Cysteine, due to its low natural frequency, exemplifies this issue. Its specific and strong chemical interactions (e.g., disulfide bonds) are observed over an extremely low random expectation, leading to statistically extreme enrichment scores (e.g., CYS-CYS value of +5.43 in the Heavy Chain). The high values of CYS in the comparison is influenced by the low abundance of Cysteine.

To investigate residue pairing preferences at interfaces, we analyzed contact frequencies and normalized contact frequencies, across antibody-antigen (Ab-Ag) complexes, protein-protein interactions (PPIs), and intra-protein contacts (Tables 10-11, Figures 8-9). To ensure our analysis of intra-protein contacts captured interactions relevant to the global protein fold, we defined a contact as a pair of spatially close residues with a minimum sequence separation of 10. This step filters out contacts between sequentially adjacent residues, which are expected to be close in space and are less informative about the protein’s tertiary structure. By focusing on these non-local interactions, we can better identify the residue pairings that stabilize the overall folded conformation. The results consistently reveal distinct patterns that differentiate Ab-Ag interfaces from general PPIs and protein interiors.

**Figure 8.**
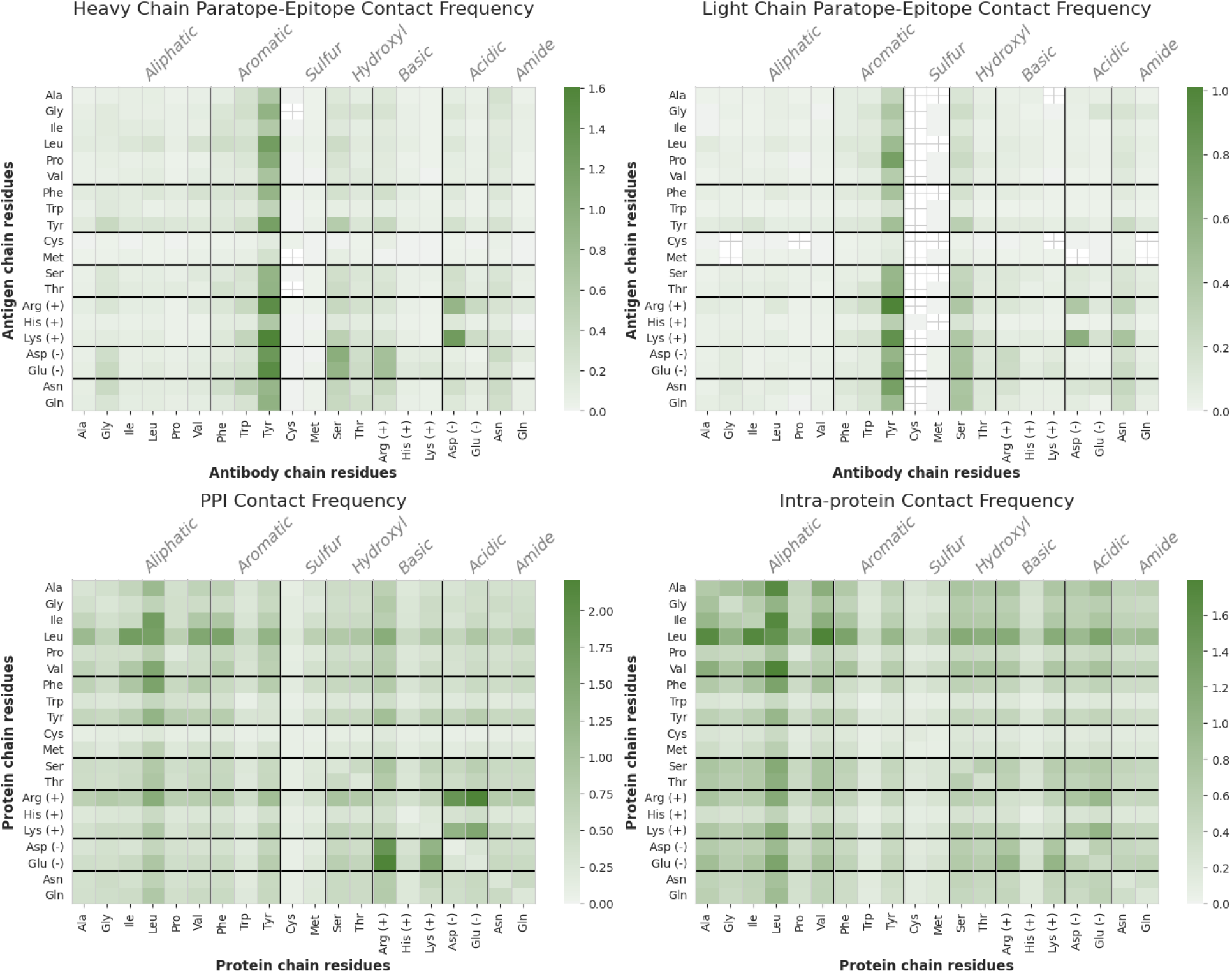
Residue pair contact frequencies in antibody-antigen (Ab-Ag) interfaces, protein-protein interactions (PPIs), and intra-protein contacts. These heatmaps illustrate the most common residue-residue pairings that define the structural architecture of different interface types. (Top Row) Antibody-antigen (Ab-Ag) interfaces are uniquely dominated by contacts involving Tyrosine (TYR), which serves as a versatile hub interacting frequently with charged, polar, and hydrophobic residues. (Bottom Left) General protein-protein (PPI) interfaces display a balanced architecture, relying on both high-frequency hydrophobic pairs (e.g., LEU-ILE) to form a core and electrostatic salt bridges (e.g., ARG-GLU) to provide specificity. (Bottom Right) Intra-protein contacts are overwhelmingly characterized by strong hydrophobic packing, with aliphatic-aliphatic interactions being the most prevalent, reflecting the drive to form a stable protein core.

**Figure 9.**
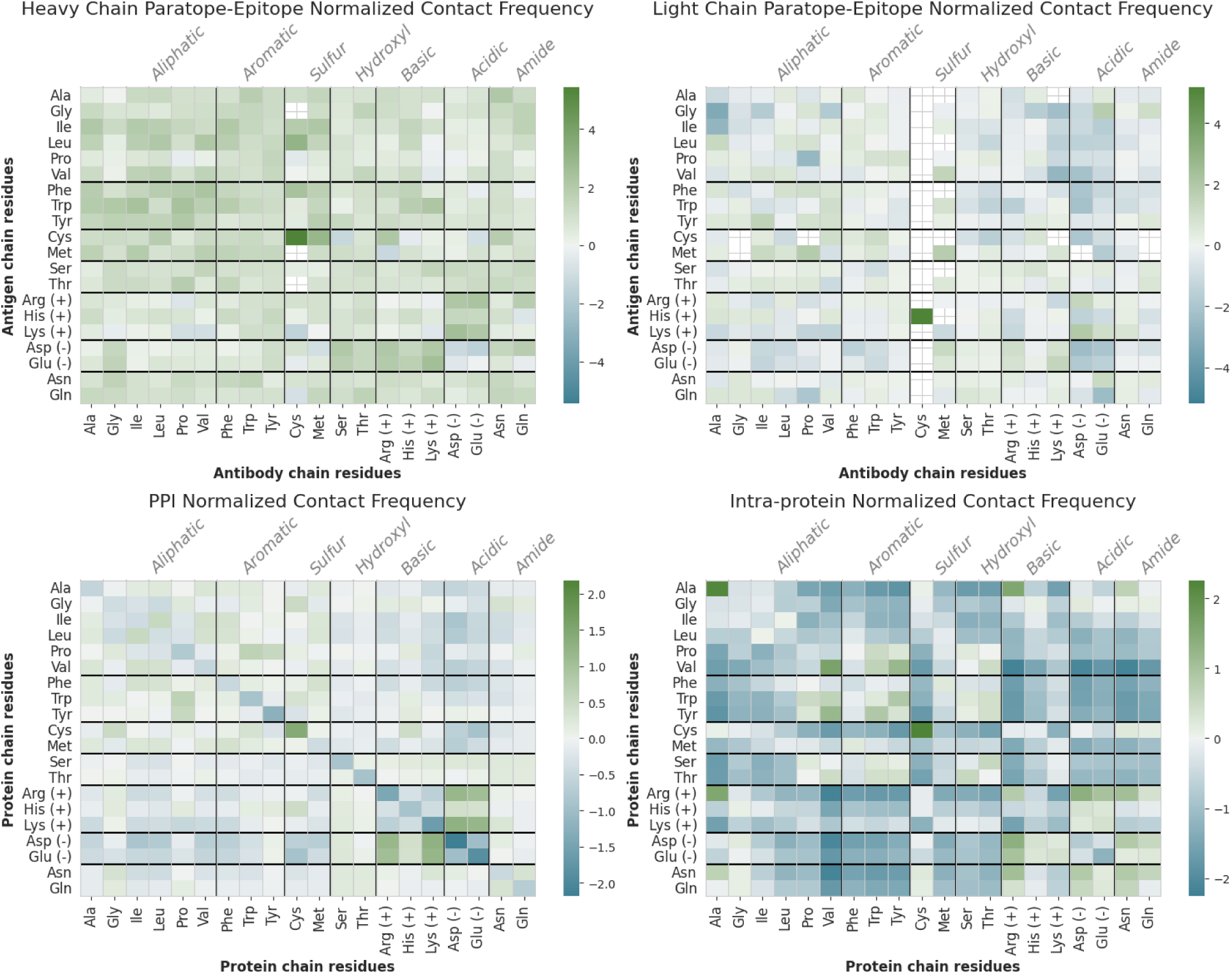
*Normalized contact frequencies of residue pairs in Ab-Ag interfaces, PPIs, and intra-protein contacts.* These heatmaps show log2 (log-odds) normalized frequencies, which correct for residue abundance to reveal underlying chemical affinities. (Top Row) Antibody-antigen interfaces show a strong intrinsic preference for interactions involving aromatic residues (TYR, TRP) and, critically, an even stronger preference for specific salt bridges (e.g., GLU-ARG, ASP-LYS) compared to general PPIs. (Bottom Left) PPI interfaces show the highest enrichment for canonical salt bridges (e.g., ARG-GLU, ARG-ASP) and are less favorable toward many aromatic pairings. (Bottom Right) Intra-protein contacts show a strong preference for CYS-CYS pairs, indicative of disulfide bonds, and disfavor many charged interactions within the hydrophobic core. The extreme positive scores for Cysteine pairs across all types reflect their structural importance and statistical rarity.

**Table 10.** Top residue-residue contact frequencies in antibody-antigen (Ab-Ag) interfaces, protein-protein interactions (PPIs), and intra-protein contacts. This table presents the most frequent residue pairings, which represent the dominant architectural “building blocks” for antibody-antigen (Ab-Ag) complexes, general protein-protein interactions (PPIs), and intra-protein contacts. The high-frequency pairs in Ab-Ag interfaces (e.g., TYR-LYS, TYR-ARG) reveal an architecture dominated by the versatile aromatic residue Tyrosine. In contrast, PPIs are built on a balanced foundation of both electrostatic salt bridges (e.g., ARG-GLU) and hydrophobic pairs (e.g., LEU-ILE), while intra-protein contacts are overwhelmingly shaped by hydrophobic pairings (e.g., LEU-ILE, LEU-VAL) that form the stable core.

| type | Heavy Chain |  | Light Chain |  | PPI |  | Intra-protein contacts |  |
| --- | --- | --- | --- | --- | --- | --- | --- | --- |
|  | aa3_aa3 | % Contact Frequency | aa3_aa3 | % Contact Frequency | aa3_aa3 | % Contact Frequency | aa3_aa3 | % Contact Frequency |
| rank |  |  |  |  |  |  |  |  |
| 1 | TYR_LYS | 4.89 | TYR_ARG | 3.08 | ARG_GLU | 2.21 | ILE_LEU | 2.57 |
| 2 | TYR_GLU | 4.52 | TYR_LYS | 2.72 | ARG_GLU | 2.21 | ILE_LEU | 2.57 |
| 3 | TYR_ARG | 4.45 | TYR_ASN | 2.30 | ARG_ASP | 1.85 | LEU_VAL | 2.56 |
| 4 | TYR_ASP | 3.99 | TYR_PRO | 2.30 | ARG_ASP | 1.85 | LEU_VAL | 2.56 |
| 5 | ASP_LYS | 3.83 | TYR_GLU | 2.03 | ILE_LEU | 1.70 | LEU_LEU | 2.20 |
| 6 | TYR_LEU | 3.77 | ASP_LYS | 1.86 | ILE_LEU | 1.70 | LEU_PHE | 2.08 |
| 7 | TYR_TYR | 3.63 | TYR_THR | 1.73 | LEU_LEU | 1.66 | LEU_PHE | 2.08 |
| 8 | TYR_PRO | 3.15 | TYR_ASP | 1.71 | LEU_PHE | 1.57 | ILE_VAL | 1.66 |
| 9 | SER_ASP | 3.03 | TYR_SER | 1.65 | LEU_PHE | 1.57 | ILE_VAL | 1.66 |
| 10 | TYR_GLN | 2.86 | TYR_LEU | 1.56 | GLU_LYS | 1.51 | ALA_LEU | 1.61 |
| 11 | TYR_GLY | 2.80 | TYR_GLN | 1.56 | GLU_LYS | 1.51 | ALA_LEU | 1.61 |
| 12 | TYR_THR | 2.78 | TYR_TYR | 1.46 | LEU_VAL | 1.51 | LEU_TYR | 1.41 |
| 13 | SER_GLU | 2.73 | TYR_PHE | 1.35 | LEU_VAL | 1.51 | LEU_TYR | 1.41 |
| 14 | TYR_SER | 2.72 | SER_GLN | 1.34 | ARG_LEU | 1.38 | ILE_PHE | 1.29 |
| 15 | TYR_PHE | 2.71 | SER_GLU | 1.34 | ARG_LEU | 1.38 | ILE_PHE | 1.29 |
| 16 | ASP_ARG | 2.68 | ASN_LYS | 1.33 | LEU_TYR | 1.25 | PHE_VAL | 1.24 |
| 17 | TYR_ASN | 2.64 | SER_ASP | 1.32 | LEU_TYR | 1.25 | PHE_VAL | 1.24 |
| 18 | ARG_GLU | 2.26 | SER_ASN | 1.30 | ASP_LYS | 1.25 | ALA_VAL | 1.16 |
| 19 | ARG_ASP | 2.23 | TYR_GLY | 1.24 | ASP_LYS | 1.25 | ALA_VAL | 1.16 |
| 20 | TYR_VAL | 2.14 | SER_ARG | 1.23 | ALA_LEU | 1.13 | ARG_LEU | 1.08 |

**Table 11.** *Top normalized contact frequencies of residue pairs in Ab-Ag interfaces, PPIs, and intra-protein contacts.* This table lists the most enriched residue pairs based on their normalized contact frequency, a score that corrects for amino acid abundance to highlight true chemical affinities. In antibody-antigen (Ab-Ag) interfaces, specific electrostatic pairs (e.g., GLU-ARG, ASP-LYS) show the strongest chemical preference, even more so than in general PPIs. This finding clarifies that antibodies utilize a multi-layered strategy combining these high-affinity electrostatic “hot spots” with a broad network of other favorable aromatic interactions. Across all interface types, Cysteine-containing pairs (e.g., CYS-CYS, CYS-HIS) exhibit exceptionally high scores, which reflect their statistical rarity and critical, non-random structural roles rather than a general chemical affinity.

| type | Heavy Chain |  | Light Chain |  | PPI |  | Intra-Protein Contacts |  |
| --- | --- | --- | --- | --- | --- | --- | --- | --- |
|  | aa3_aa3 | Normalized Contact Frequency | aa3_aa3 | Normalized Contact Frequency | aa3_aa3 | Normalized Contact Frequency | aa3_aa3 | Normalized Contact Frequency |
| rank |  |  |  |  |  |  |  |  |
| 1 | CYS_CYS | 5.43 | CYS_HIS | 5.18 | CYS_CYS | 1.42 | CYS_CYS | 1.71 |
| 2 | CYS_LEU | 3.09 | ASP_LYS | 1.92 | ASP_LYS | 1.29 | GLU_LYS | 1.16 |
| 3 | MET_CYS | 2.9 | PRO_MET | 1.84 | ASP_LYS | 1.29 | GLU_LYS | 1.16 |
| 4 | ASP_LYS | 2.59 | MET_MET | 1.81 | GLU_LYS | 1.19 | ASP_LYS | 1.13 |
| 5 | LYS_GLU | 2.58 | GLU_GLY | 1.78 | GLU_LYS | 1.19 | ASP_LYS | 1.13 |
| 6 | ILE_TRP | 2.4 | ILE_TYR | 1.53 | ARG ASP | 1.17 | ARG_GLU | 1.12 |
| 7 | PRO_TRP | 2.39 | MET_VAL | 1.42 | ARG ASP | 1.17 | ARG_GLU | 1.12 |
| 8 | CYS_PHE | 2.38 | LYS_GLU | 1.38 | ARG_GLU | 1.05 | ARG ASP | 1.12 |
| 9 | GLU_ARG | 2.3 | ASP_ARG | 1.36 | ARG_GLU | 1.05 | ARG ASP | 1.12 |
| 10 | VAL_PHE | 2.27 | ALA_LEU | 1.22 | PRO_TRP | 0.65 | ASP_HIS | 0.59 |
| 11 | LYS_TRP | 2.2 | ILE_MET | 1.21 | PRO_TRP | 0.65 | ASP_HIS | 0.59 |
| 12 | MET_ILE | 2.16 | MET ASP | 1.2 | PRO_TYR | 0.54 | PRO_TRP | 0.54 |
| 13 | ARG_CYS | 2.14 | GLU ASN | 1.18 | PRO_TYR | 0.54 | PRO_TRP | 0.54 |
| 14 | VAL_LEU | 2.13 | TRP_CYS | 1.08 | ILE_LEU | 0.52 | ASN ASP | 0.5 |
| 15 | ASP_ARG | 2.13 | GLN_GLY | 1.04 | ILE_LEU | 0.52 | ASN ASP | 0.5 |
| 16 | LEU_LEU | 2.12 | VAL_CYS | 1.01 | CYS_GLY | 0.48 | ASN_SER | 0.41 |
| 17 | ALA_ILE | 2.11 | PRO_PHE | 1 | CYS_GLY | 0.48 | ASN_SER | 0.41 |
| 18 | GLU_LYS | 2.1 | GLU_LYS | 0.99 | CYS_HIS | 0.46 | PRO_TYR | 0.38 |
| 19 | ASN_ALA | 2.09 | VAL_TRP | 0.99 | CYS_HIS | 0.46 | PRO_TYR | 0.38 |
| 20 | ARG ASP | 2.08 | ARG ASP | 0.99 | ASP_HIS | 0.41 | GLU_HIS | 0.38 |

In antibody-antigen interfaces, contacts involving aromatic residues, particularly Tyrosine, are the most frequent. These pairings, including aromatic-charged (e.g., TYR-LYS) and aromatic-acidic (e.g., TYR-GLU, TYR-ASP), rank among the most common in heavy and light chain paratope-epitope contacts, reflecting the well-established role of tyrosine in mediating versatile binding through hydrophobic packing, hydrogen bonding, and cation–π interactions (Kunik and Ofran 2013; Reis et al. 2022). In contrast, general PPI interfaces are dominated by a combination of classical salt bridges (ARG-GLU, ARG-ASP) and hydrophobic pairs (LEU-ILE, LEU-LEU), which are less frequent in antibody binding, emphasizing a shift toward aromatic-driven interactions in immune recognition (Table 10).

Normalized contact frequencies (Table 11) highlight the intrinsic chemical preferences, correcting for residue abundance, and reveal a more nuanced binding strategy for antibodies. While aromatic pairs are broadly favored, the analysis shows that the strongest chemical preferences in the heavy chain are for specific electrostatic pairs. For example, the normalized score for GLU-ARG is +2.30 and for ASP-LYS is +2.59, placing them among the most enriched interactions. Crucially, this contradicts the narrative that antibodies de-emphasize salt bridges; in fact, the chemical preference for these pairs is stronger in antibodies than in general PPIs. The ARG-ASP pair has a normalized score of +2.08 in the heavy chain, nearly double its score of +1.17 in PPIs, indicating a much stronger intrinsic preference in the antibody interface. Furthermore, despite cysteine’s overall rarity, CYS-containing pairs exhibit remarkable outlier specificities. CYS-CYS contacts, crucial for forming disulfide bonds, show exceptional enrichment in the Heavy Chain (+5.43), and the CYS-HIS contact is the most enriched pair in the Light Chain (+5.18). These findings collectively underscore that the antibody binding strategy is a sophisticated expansion of the chemical repertoire, combining a high-frequency scaffold of versatile aromatic contacts with less frequent but more strongly preferred electrostatic “hot spots” to achieve both specificity and affinity.

Overall, these findings align with previous studies showing that antibody binding sites are compositionally distinct (Kunik and Ofran 2013; Reis et al. 2022; Nadalin and Carbone 2018). However, our analysis clarifies that compared to PPIs, which rely on a balance of frequent hydrophobic and electrostatic contacts, antibodies employ a multi-layered strategy. They leverage a chemically diverse interface, built upon a high-frequency aromatic network, which is then augmented by exceptionally strong chemical preferences for specific salt bridges and structurally critical cysteine linkages. This unique combination enables the flexible recognition of highly variable antigens, underscoring their evolutionary adaptation for a broad and specific immune response.

